# AGNOSTOS-DB: a resource to unlock the uncharted regions of the coding sequence space

**DOI:** 10.1101/2021.06.07.447314

**Authors:** Chiara Vanni, Matthew S. Schechter, Tom O. Delmont, A. Murat Eren, Martin Steinegger, Frank Oliver Glöckner, Antonio Fernandez-Guerra

## Abstract

Genomes and metagenomes contain a considerable percentage of genes of unknown function, which are often excluded from downstream analyses limiting our understanding of the studied biological systems. To address this challenge, we developed AGNOSTOS, a combined database-computational workflow resource that unifies the known and unknown coding sequence space of genomes and metagenomes. Here, we present AGNOSTOS-DB, an extensive database of high-quality gene clusters enriched with functional, ecological and phylogenetic information. Moreover, AGNOSTOS allows integrating new data into existing AGNOSTOS-DBs, maximizing the information retrievable for the genes of unknown function. As a proof of concept, we provide a seed database that integrates the predicted genes from marine and human metagenomes, as well as from Bacteria, Archaea, Eukarya and giant viruses environmental and cultivar genomes. The seed database comprises 6,572,081 gene clusters connecting 342 million genes and represents a comprehensive and scalable resource for the inclusion and exploration of the unknown fraction of genomes and metagenomes.

## Background & Summary

The characterization of genes of unknown function can catalyze research in basic science, medicine, and biotechnology^1^. Approximately 30-35% of the genes in genomes and metagenomes lack functional characterization^2^. The constant influx of newly sequenced microbial and viral genomes through cultivation, single-cell genomics and metagenomics is increasing the volume of genes of unknown function. Recently we developed AGNOSTOS^2^, a conceptual framework and a computational workflow that unifies the known and the unknown coding sequence space (CDS-space) of genomes and metagenomes to address the limitations of the current approaches. Such limitations include the lack of a systematic categorization of the genes with unknown function into biologically meaningful categories, the inclusion of both genomic and metagenomic data and often an overestimation of the number of unknowns due to the lack of remote-homology searches. AGNOSTOS partitions the CDS-space using gene clusters, and unlike the previous GC centered methods^3–6^, performs an extensive validation yielding high-quality biologically and phylogenetically aware functional units. Further, AGNOSTOS provides a thorough characterization of the unknown space, including distant-homology classification methods. Moreover, GCs sharing remote homologies are aggregated in communities (GCCs)^2^. Altogether AGNOSTOS allows exploring the CDS-space at different levels: from the gene similarities within a GC to the remote homologies in the GCCs.

The AGNOSTOS databases (AGNOSTOS-DBs) are embedded into the AGNOSTOS analytical environment and allow different levels of analysis, as shown in Figure 1. The AGNOSTOS-DB provides GC sequence profiles that can be used to quickly screen new datasets (*profile-search* module). Alternatively, a novel DB can be created from any metagenomic and genomic data via the *DB_creation* module. Finally, a key feature of AGNOSTOS is its scalability, i.e., the possibility to update existing DBs with new sequences.

**Figure 1.**
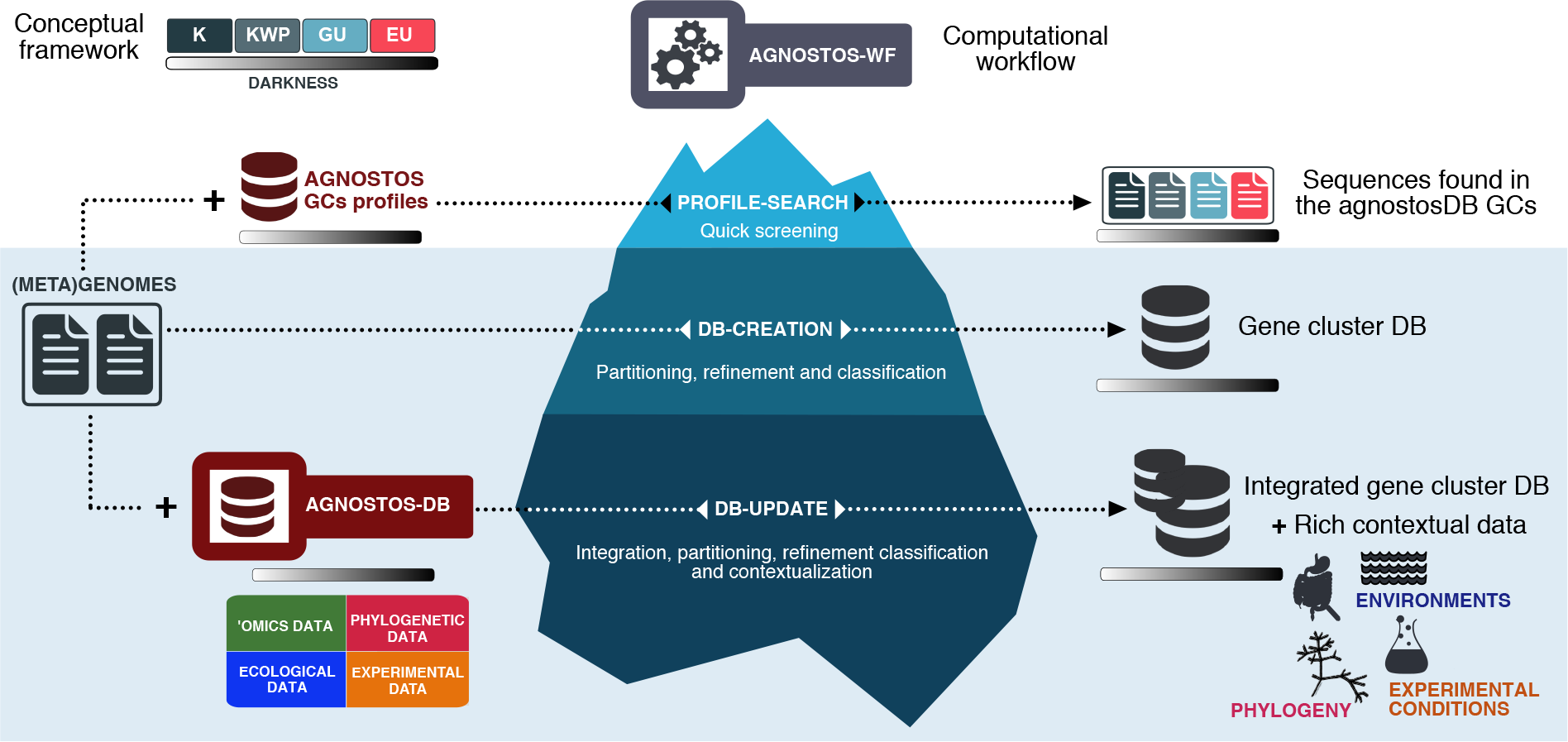
Schematic overview of the AGNOSTOS analytical environment (workflow and databases).

The AGNOSTOS seed-DB integrated the sequence data from 1,829 marine and human metagenomic assemblies and 28,941 bacterial and archaeal genomes, for a total of 5,287,759 GCs and 335,439,673 genes^2^. To show the flexibility and scalability provided by AGNOSTOS, here we expanded the seed-DB by integrating genomes affiliated to Eukarya and their infecting nucleocytoplasmic large DNA viruses (NCLDV) characterized mostly from metagenomic sequence data corresponding to the surface of the oceans (Gaia et al., personal communication). First, we integrated the predicted genes from 3,243 NCLDV environmental and cultivar genomes^7–9^<u>(Gaia et al., personal communication)</u>, obtaining a database that contains 5,383,876 GCs and 336,513,365 genes (Table 1).

**Table 1.** Seed + NCLDV database. Number of genes, GCs, and GCCs per category.

|  | K | KWP | GU | EU | Total |
| --- | --- | --- | --- | --- | --- |
| Genes | 230,972,742 | 32,974,245 | 68,898,794 | 3,667,584 | 336,513,365 |
| GCs | 1,680,863 | 799,647 | 2,678,313 | 225,053 | 5,383,876 |
| GCCs | 75,332 | 119,111 | 443,961 | 122,812 | 761,216 |

Subsequently, we integrated the predicted genes from 713 eukaryotic environmental genomes^10^, creating a comprehensive database of 6,572,081 GCs and 341,655,294 genes (Table 2) that expand to all domains of life and viruses.

**Table 2.**
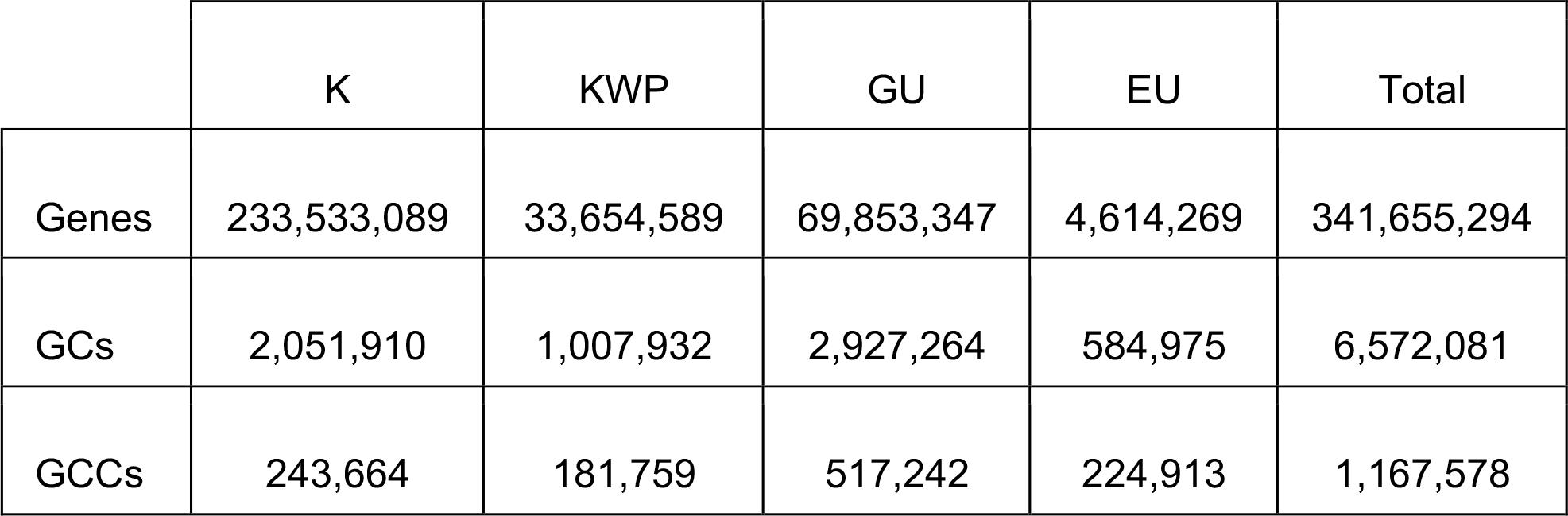
AGNOSTOS seed + NCLDV + Eukaryotic plankton GC database. Number of genes, GCs and GCCs by category.

|  | K | KWP | GU | EU | Total |
| --- | --- | --- | --- | --- | --- |
| Genes | 233,533,089 | 33,654,589 | 69,853,347 | 4,614,269 | 341,655,294 |
| GCs | 2,051,910 | 1,007,932 | 2,927,264 | 584,975 | 6,572,081 |
| GCCs | 243,664 | 181,759 | 517,242 | 224,913 | 1,167,578 |

The AGNOSTOS GCs proved to be relatively stable after integrating new data. The GCs updated with new genes showed a decrease in the intra-cluster average similarity^2^ of 7% after the first integration and 11% after the second. The overall intra-cluster average similarity values decreased by 1% with the two integrations (Table 3).

**Table 3.**
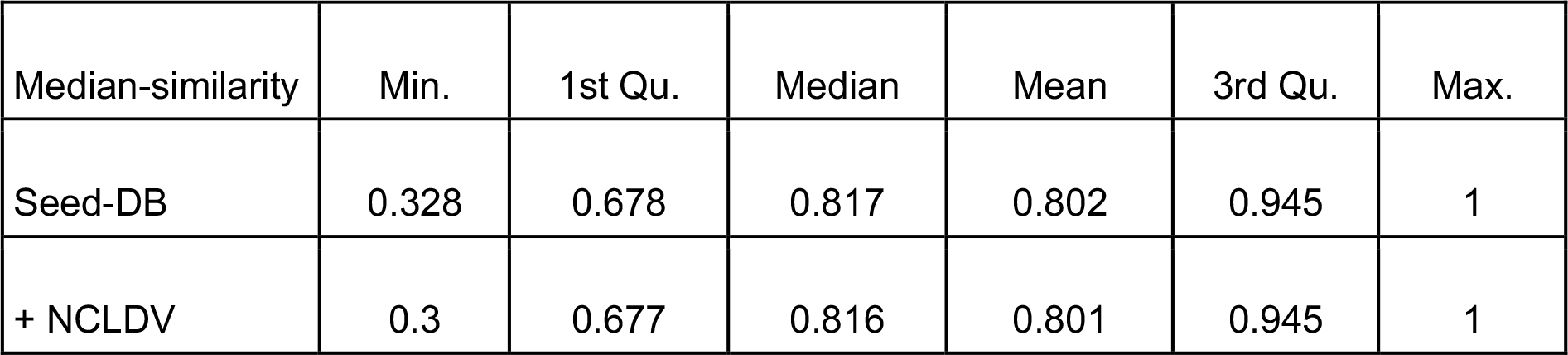
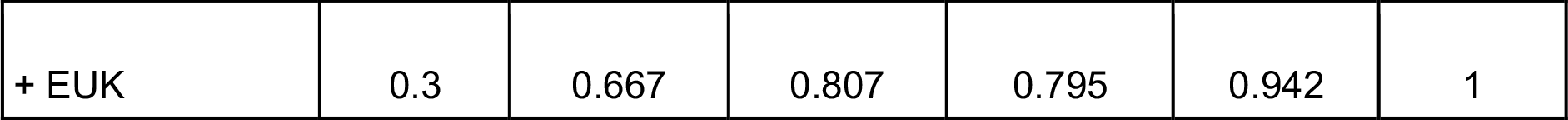
Changes in the GCs median similarity values after the integration of NCLDV and plankton eukaryotic genes. (The values are filtered for clusters with more than 2 members.)

| Median-similarity | Min. | 1st Qu. | Median | Mean | 3rd Qu. | Max. |
| --- | --- | --- | --- | --- | --- | --- |
| Seed-DB | 0.328 | 0.678 | 0.817 | 0.802 | 0.945 | 1 |
| + NCLDV | 0.3 | 0.677 | 0.816 | 0.801 | 0.945 | 1 |
| + EUK | 0.3 | 0.667 | 0.807 | 0.795 | 0.942 | 1 |

In general, the integration of new data led to the reduction of the original set of singletons (Table 4), suggesting that many of these sequences are not just artifacts or the results of sequencing and assembly errors. However, 241,669 NCLDV and 4,264,489 eukaryotic genes were added to the singletons set (Table 4), increasing the total number of singletons by 17%, manifesting the novelty added when we integrate poorly characterized and phylogenetically diverse groups such as the NCLDVs and the plankton eukaryotes.

**Table 4.** Changes in the number of singletons as a consequence of the integration of NCLDVs and plankton eukaryotic sequences.

| Seed-DB | Seed-DB+NCLDV | Seed-DB+NCLDV+EUK |
| --- | --- | --- |
| 24,977,524 | 25,181,853 | 29,328,400 |
| Seed-DB |  |  |
| 24,977,524 | Seed-DB+NCLDV |  |
| - 37,340<br>+ 241,669 = | 25,181,853 | Seed-DB+NCLDV+EUK |
|  | - 117,942<br>+ 4,264,489 = | 29,328,400 |

Moreover, after the two integrations, we observed an increase of 23% in unknown GCs (187% for the EU GCs alone) compared to the seed-DB. These novel GCs found in the context of NCLDV and eukaryotic genomes offer a starting point for targeted analyses of the unknown fraction. Ultimately, as shown in Figure 2, we found most GCs unique to the prokaryotic or eukaryotic plankton fraction where more than half of the GCs in these two sets belong to the unknown fraction (Figure 2). We also identified 5,220 GCs shared by all datasets and 2,854 GCs shared between the NCLDVs and the plankton eukaryotes, in both cases mainly members of the known fraction (Figure 2). This last set constitutes a valuable resource to better understand the relationship between the NCLDVs and their eukaryotic hosts^11^.

**Figure 2.**
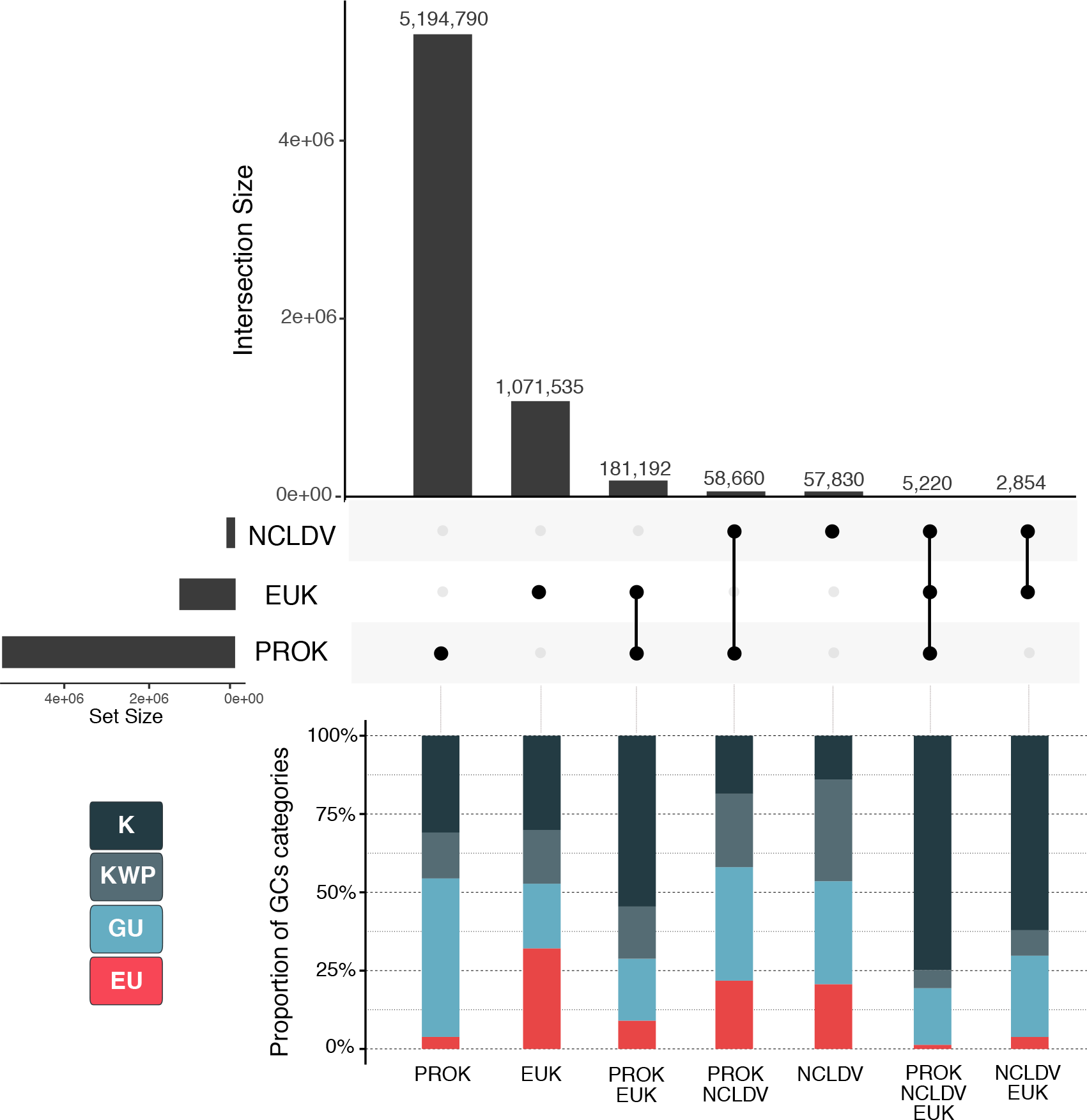
GC distribution in the different datasets and proportion of categories per set. NCLDV=giant virus GCs, EUK=plankton eukaryotic GCs and PROK=seed-DB archaea and bacterial GCs.

The seed-DB and the databases resulting from the integration of NCLDV and plankton eukaryotes genomes are publicly available on Figshare (Data Citation 1-3). The workflow code is available on GitHub (https://github.com/functional-dark-side/agnostos-wf).

## Methods

### Computational workflow

We implemented the AGNOSTOS using the Snakemake workflow management system^12^ to easily process large datasets in a reproducible manner. The workflow is publicly available in a GitHub repository at https://github.com/functional-dark-side/agnostos-wf (version 1.0, used in this paper, is archived at https://doi.org/10.5281/zenodo.4557847). The workflow provides three different modules (Fig. 1): The *DB-creation* module, which creates a de-novo GC dataset, taking as input metagenomic contigs or predicted genes in *fastA* format. The output is a dataset of characterized GCs, partitioned in four categories (Fig. 1). The output of the *DB-creation* module is already formatted to be updated with new data via the *DB-update* module. The *DB-update* module allows the continuous integration of new sequence data into an existing GC database. The new sequences are integrated into existing clusters if similar enough or clustered in new GCs if not. The final output consists of the updated GC database and related files containing contextual data about the GCs.

Moreover, the workflow provides a *profile-search* module that enables the screening of existing GC databases using sensitive profile-search methods. The sequence-profile search is performed using the *search* program of the MMseqs2^13^ software (parameters: *-e 1e-20 -- cov-mode 2 -c 0*.*6*). This module takes in input the GC MMseqs2 profiles, a tabular file with the information about the GC functional categories and the sequences to search against the profiles, in *fastA* format. The search results are then filtered to keep hits within the 90% of the best *log(e-value)* and retrieve the queries consensus GC annotation via a quorum majority voting approach.

The required software tools and programs are installed via Conda or via the provided installation script. When the existing database is the seed-DB, or an updated GC database originated from the seed-DB, the *DB-update* final rule links the new integrated GCs with the ecological, phylogenetic, and experimental metadata obtained from the Vanni et al. analyses^2^ (Fig. 1).

To build the GC databases presented here, we ran the workflow in the de.NBI Cloud (https://www.denbi.de/cloud)^14^, using a cluster setup with 10 nodes of 28 cores and 252G of memory each. The cluster was built using BiBiGrid (https://github.com/BiBiServ/bibigrid) and it is using SLURM^15^ for job scheduling and cluster management.

### Data collection

To build the seed-DB, we combined a set of 1,829 metagenomes from five major metagenomic surveys of the ocean and human microbiome with 28,941 archaea and bacterial genomes^2^. The data sources per project are specified in Table 5-A, and the number of genes predicted from each project in Table 5-B. The metagenomic contextual data is found in one SQLite (version 3.25.0) (https://www.sqlite.org/index.html) database “contextual_data.db”, which is available in the “agnostosDB_dbf02445-20200519_environmental” folder (download: https://ndownloader.figshare.com/files/23066879) within the seed-DB dataset deposited in Figshare (https://doi.org/10.6084/m9.figshare.12459056, Data Citation 1)

**Table 5.** AGNOSTOS seed-DB source datasets.

(A) Metagenomic and genomic project data sources.
| Dataset | Reference | Raw data | Contextual data |
| --- | --- | --- | --- |
| TARA | Sunagawa et al.<sup>16</sup> | ENA | PANGAEA |
| Malaspina | Duarte et al.<sup>17</sup> | Duarte et al.<sup>17</sup> | Duarte et al.<sup>17</sup> |
| OSD | Kopf et al.<sup>18</sup> | ENA | PANGAEA |
| GOS | Rush et al.<sup>19</sup> | NCBI | iMICROBE |
| HMP | Lloyd-Price et al.<sup>20</sup> | HMP portal | HMP portal |
| GTDB | Parks et al.<sup>21</sup> | Annotree<sup>22</sup> | Annotree<sup>22</sup> |

(B) Number of genomes, metagenomes and predicted genes per project.
| Dataset | (Meta)genomes | Predicted genes |
| --- | --- | --- |
| TARA | 242 | 111,903,261 |
| Malaspina | 116 | 20,574,033 |
| OSD | 145 | 7,015,383 |
| GOS | 80 | 20,068,580 |
| HMP | 1,246 | 162,687,295 |
| GTDB-r86 | 28,941 | 93,723,190 |

We expanded the seed-DB by integrating 3,243 cultivar and environmental genomes from nucleocytoplasmic large DNA viruses (NCLDVs). This collection combines the environmental genomes generated by Moniruzzaman et al.^7^, Schultz et al.^8^, and by the TARA Oceans Consortium (Gaia et al., personal communication, dataset unpublished) as well as 235 reference genomes collected from GenBank^9^ (Table 6).

**Table 6.** The NCLDV genomes dataset

| Dataset | Genomes | Predicted genes |
| --- | --- | --- |
| Moniruzzaman | 501 | 631,572 |
| Schultz | 1,762 | 181,068 |
| TARA | 745 | 193,855 |
| GenBank | 235 | 99,599 |

The NCLDV dataset was passed to AGNOSTOS in the form of predicted genes (amino acids).

Additionally, we added to the seed+NCLDV-DB, 10,207,450 eukaryotic plankton predicted genes, obtained from the combination of 683 environmental genomes and 30 single amplified genomes^10^. The plankton eukaryotic genes were predicted by Delmont et al. using a combination of three approaches: a homology-based search against protein reference databases (Uniref90 + METdb), the mapping of the metatranscriptomics assemblies to the environmental and single-amplified genomes, and an ab-initio gene prediction using *gmove* (http://www.genoscope.cns.fr/externe/gmove/) ^10^.

Both database integrations were performed using the *DB-update* module of the workflow.

### Update of a gene cluster dataset

AGNOSTOS updates an existing GC database using the *clusterupdate* module of MMseqs2^23^ using the same clustering parameters described in Vanni et al.^2^. The output of the clustering update is then parsed to retrieve three GC datasets: (1) The “original” GCs, a set containing the original GCs without any similarity to the new sequences. (2) The “shared” set of GCs with mixed, original and new, sequences. (3) The “new” GCs built from the new sequences without any similarity to the existing GCs. The “new” GCs and the shared GCs that were originally singletons, are processed through all the workflow steps until the GCC inference.

*Results of the integration and gene cluster contextualization*. At the end of the *DB-update* module, the three GC sets, together with their additional information, are re-combined in a single GC dataset. In case the original GC database originated from the seed-DB^2^, the final GC dataset can be decorated with the ecological and phylogenetic contextual data described in Vanni et al.^2^. This information is included in the online version of the seed-DB (Data Citation 1), and can be used at the end of the *DB-update* module to further characterize the updated GCs (Fig. 1).

## Data Records

The AGNOSTOS seed-DB is available on Figshare at https://doi.org/10.6084/m9.figshare.12459056 (Data Citation 1). We base encoded the seed version of the database as agnostosDB_dbf02445-20200519, which resulted from using the *crc32* hash algorithm to encode the word “seed”, followed by the release date (YYYYMMDD). Future integrations of (meta)genomic data into the seed-DB will be named by adding the new dataset name in the *crc32* encoding command followed by the new release date. The seed-DB is a dataset of microbial GCs containing 40,423,549 GCs with 415,971,742 genes predicted from 28,941 bacterial and archaeal genomes and 1,749 metagenomes.

The “agnostosDB_dbf02445-20200519_original-data” (https://ndownloader.figshare.com/files/23066858) contains the list of metagenomes, genomes, contigs and predicted genes per metagenomic project and found in GTDB. The original seed-DB GC MMseqs2 database files are available in “agnostosDB_dbf02445-20200519_mmseqs_clustering” (https://ndownloader.figshare.com/files/23066651). The distribution of the GCs and genes in the four categories can be found in Supplementary Figure 2 of Vanni et al.^2^. The list of refined GCs, their categories and their genes are available in the TSV file: “cluster_ids_categ_genes.tsv.gz”. The TSV file “HQ-clusters.tsv.gz” contains the IDs of the set of high-quality GCs, containing mostly complete genes. The GC HH-suite database files are contained in the folder “agnostosDB_dbf02445-20200519_hh-suite-db” (https://ndownloader.figshare.com/files/23064476). The HH-suite database can be used to screen the GCs in search for remote-homologies, using HHblits^24^ (see Usage notes). The GC MMseqs2 profiles used by the *profile-search* module are stored in the folder “agnostosDB_dbf02445-20200519_mmseqs_profiles” (https://ndownloader.figshare.com/files/23066963).

All other files needed by the *DB-update* and the *profile-search* modules are also available in the database Figshare folder.

The GC database expanded with the NCLDV MAGs was encoded as agnostosDB_a42ac58a-20200715 and it is available on Figshare at https://doi.org/10.6084/m9.figshare.13251743 (Data citation 2).

The database agnostosDB_4eab867d-20201104 combining the seed-DB GCs with the NCLDV and the eukaryotic plankton GCs is available on Figshare at https://doi.org/10.6084/m9.figshare.13264769 (Data citation 3).

The file organization of the two updated GCs datasets mirrors that of the seed-DB.

## Technical Validation

### Gene cluster database validations

The quality control steps in AGNOSTOS thoroughly validate the AGNOSTOS-DB GCs. These steps include the identification and removal of eventual spurious genes, the validation of the GCs internal homogeneity, both in terms of Pfam protein domain annotations and sequence composition, and a remote-homology refinement of the GCs classification in categories, which aimed to avoid an overestimation of the real number of uncharacterized genes^2,25^.

### Computational workflow validations

We followed the FAIR guiding principles^26^ to create a reproducible and reusable workflow. We implemented AGNOSTOS using the Snakemake workflow management system^12^ and is available as an open source software (https://doi.org/10.5281/zenodo.4557847). The majority of the dependencies can be installed within a Conda environment^27,28^ integrated in the workflow. We provide an installation script to install the software not available in Conda. The workflow can be tested using the provided test datasets for the *DB-creation* and *DB-update* modules of AGNOSTOS. The test datasets are intended to validate the installation and help the user with an easy example of the workflow functioning and extensive outputs. For the testing dataset, we selected three TARA Ocean surface samples from the Indian Ocean (IO) (TARA_038, TARA_039 and TARA_041), all within a size fraction of 0.1-0.22 micro-m (giant viruses, archaea and bacteria enriched fraction) and we randomly chose 5K contigs from each sample. The test dataset is available for download from Figshare (https://doi.org/10.6084/m9.figshare.12630581, Data Citation 4).

## Usage Notes

The workflow is accompanied by detailed usage instructions and description of the output files available in the GitHub repository (https://github.com/functional-dark-side/agnostos-wf).

The GC profiles and the tables with the GC categories used by the *profile-search* module can be downloaded from their respective AGNOSTOS-DB Figshare folders.

Any other MMseqs2 profile database can be used for the search. For example, to screen only the seed-DB high-quality GCs, we can use their identifiers from the TSV file “HQ_clusters.tsv.gz”, to subset the full GC profile database with the MMseqs2 *createsubdb* module (see the MMseqs2 user guide available at https://mmseqs.com/latest/userguide.pdf).

By default, the *DB-update* module of the workflow will download and use the seed-DB as the existing GC database. Any other GC database built with the *DB-creation* module, or containing the same output file structure, can be used by specifying the different database location in the *DB-update* configuration file.

The output of the *DB-update* and *DB-creation* modules can be analyzed using anvi’o^29^, by integrating all the information gathered for the GCs as “functions” in the anvi’o contig database. An introduction on how to integrate AGNOSTOS products in the analyses with anvi’o is available in the form of a tutorial at https://merenlab.org/agnostos-tutorial. The tutorial describes the integration of the infant gut dataset (IGD) from Sharon et al.^30^ in AGNOSTOS (workflow and databases) and how to use it in combination with anvi’o to explore genes of unknown function. Within anvi’o, the GCs can be explored in the context of assembled contigs allowing the identification of contigs enriched in uncharacterized genes or the exploration of the genomic context of GCs of interest. Furthermore, the AGNOSTOS GCs can also be added as “functions” to the external and internal genomes storages used by anvi’o to calculate a pangenome. The pangenome can then be investigated to identify core and genome-specific GCs.

## Code Availability

The workflow is publicly available and maintained on a GitHub repository (https://github.com/functional-dark-side/agnostos-wf). The workflow has been tested in an HPC cluster setup with at least 4 nodes of 28 cores and 252 G of memory each, which uses SLURM^15^ as Grid Batch Scheduler. The workflow programs and software contained in the Conda environment are listed in a .yml file at https://github.com/functional-dark-side/agnostos-wf/blob/master/envs/workflow.yml. The other programs are listed in the bash installation script at https://github.com/functional-dark-side/agnostos-wf/blob/master/installation_script.sh.

## Notes

https://github.com/functional-dark-side/agnostos-wf

https://doi.org/10.6084/m9.figshare.13251743

https://doi.org/10.6084/m9.figshare.13264769

### Competing Interest Statement

The authors have declared no competing interest.

## Data citation

1. Vanni, Chiara; Fernandez-Guerra, Antonio (2020): agnostosDB_dbf02445-20200519. figshare. Dataset. https://doi.org/10.6084/m9.figshare.12459056

2. Vanni, Chiara; Fernandez-Guerra, Antonio (2020): agnostosDB_a42ac58a-20200715. figshare. Dataset. https://doi.org/10.6084/m9.figshare.13251743

3. Vanni, Chiara; Fernandez-Guerra, Antonio (2020): agnostosDB_4eab867d-20201104. figshare. Dataset. https://doi.org/10.6084/m9.figshare.13264769

4. Vanni, Chiara (2020): agnostos-wf test dataset. figshare. Dataset. https://doi.org/10.6084/m9.figshare.12630581

## References

1. Zhang, X. et al.. Assignment of function to a domain of unknown function: DUF1537 is a new kinase family in catabolic pathways for acid sugars. Proc. Natl. Acad. Sci. U. S. A. 113, E4161–9 (2016).

2. Vanni, C. et al.. Light into the darkness: Unifying the known and unknown coding sequence space in microbiome analyses. bioRxiv 2020.06.30.180448 (2020) doi:10.1101/2020.06.30.180448.

3. Yooseph, S. et al.. The Sorcerer II Global Ocean Sampling Expedition: Expanding the Universe of Protein Families. PLoS Biol. 5, 1–35 (2007).

4. Hurwitz, B. L. & Sullivan, M. B. The Pacific Ocean Virome (POV): A Marine Viral Metagenomic Dataset and Associated Protein Clusters for Quantitative Viral Ecology. PLoS One 8, (2013).

5. Brum, J. R. et al.. Illuminating structural proteins in viral “dark matter” with metaproteomics. Proc. Natl. Acad. Sci. U. S. A. 113, 2436–2441 (2016).

6. Wyman, S. K., Avila-Herrera, A., Nayfach, S. & Pollard, K. S. A most wanted list of conserved microbial protein families with no known domains. PLoS One 13, e0205749 (2018).

7. Moniruzzaman, M., Martinez-Gutierrez, C. A., Weinheimer, A. R. & Aylward, F. O. Dynamic genome evolution and complex virocell metabolism of globally-distributed giant viruses. Nat. Commun. 11, 1710 (2020).

8. Schulz, F. et al.. Giant virus diversity and host interactions through global metagenomics. Nature 578, 432–436 (2020).

9. Dennis Benson, G. A., Karsch-Mizrachi, I., Lipman, D. J., Ostell, J. & Wheeler, D. L. GenBank. Nucleic Acids Res. 36, 25–30 (2008).

10. Delmont, T. O. et al.. Functional repertoire convergence of distantly related eukaryotic plankton lineages revealed by genome-resolved metagenomics. (2020) doi:10.1101/2020.10.15.341214.

11. Hingamp, P. et al.. Exploring nucleo-cytoplasmic large DNA viruses in Tara Oceans microbial metagenomes. ISME J. 7, 1678–1695 (2013).

12. Köster, J. Reproducible data analysis with Snakemake. F1000Res. 7, (2018).

13. Steinegger, M. & Söding, J. Clustering huge protein sequence sets in linear time. Nat. Commun. 9, 2542 (2018).

14. Belmann, P. et al.. de.NBI Cloud federation through ELIXIR AAI. F1000Res. 8, 842 (2019).

15. Yoo, A. B., Jette, M. A. & Grondona, M. SLURM: Simple Linux Utility for Resource Management. Job Scheduling Strategies for Parallel Processing 44–60 (2003) doi:10.1007/10968987_3.

16. Sunagawa, S. et al.. Ocean plankton. Structure and function of the global ocean microbiome. Science 348, 1261359 (2015).

17. Duarte, C. M. Seafaring in the 21St Century: The Malaspina 2010 Circumnavigation Expedition. Limnol. Oceanog. Bull. 24, 11–14 (2015).

18. Kopf, A. et al.. The ocean sampling day consortium. Gigascience 4, 27 (2015).

19. Rusch, D. B. et al.. The Sorcerer II Global Ocean Sampling Expedition: Northwest Atlantic through Eastern Tropical Pacific. PLoS Biol. 5, 1–34 (2007).

20. Lloyd-Price, J. et al.. Strains, functions and dynamics in the expanded Human Microbiome Project. Nature 550, 61–66 (2017).

21. Parks, D. H. et al.. A standardized bacterial taxonomy based on genome phylogeny substantially revises the tree of life. Nat. Biotechnol. 36, (2018).

22. Mendler, K. et al.. AnnoTree: visualization and exploration of a functionally annotated microbial tree of life. Nucleic Acids Res. 47, 4442–4448 (2019).

23. Steinegger, M. & Soding, J. MMseqs2 enables sensitive protein sequence searching for the analysis of massive data sets. Nat. Biotechnol. advance on, (2017).

24. Remmert, M., Biegert, A., Hauser, A. & Söding, J. HHblits: Lightning-fast iterative protein sequence searching by HMM-HMM alignment. Nat. Methods 9, 173–175 (2012).

25. Jaroszewski, L. et al.. Exploration of Uncharted Regions of the Protein Universe. PLoS Biol. 7, e1000205–e1000205 (2009).

26. Wilkinson, M. D. et al.. The FAIR Guiding Principles for scientific data management and stewardship. Sci Data 3, 160018 (2016).

27. conda-forge community. The conda-forge Project: Community-based Software Distribution Built on the conda Package Format and Ecosystem. (2015). doi:10.5281/zenodo.4774217.

28. Grüning, B. et al.. Bioconda: sustainable and comprehensive software distribution for the life sciences. Nat. Methods 15, 475–476 (2018).

29. Eren, A. M. et al.. Community-led, integrated, reproducible multi-omics with anvi’o. Nat Microbiol 6, 3–6 (2021).

30. Sharon, I. et al.. Time series community genomics analysis reveals rapid shifts in bacterial species, strains, and phage during infant gut colonization. Genome Res. 23, 111–120 (2013).

